# High-throughput RNA and Single-cell RNA Sequencing to Identify Sepsis associated Molecular Mechanisms and the Blood Immune Cell Landscape

**DOI:** 10.1101/2025.03.31.646322

**Authors:** Liu Wenjun, Dezhong Cheng

## Abstract

Sepsis, a leading cause of death in intensive care units (ICUs), is a complex systemic inflammatory response to infection with high morbidity and mortality. Its pathogenesis involves dysregulated inflammation, immune dysfunction, and metabolic alterations, particularly in lactate metabolism. This study employed bioinformatics analyses to explore sepsis mechanisms and identify potential therapeutic targets.

We analyzed two GEO datasets and found the lactate metabolism pathway significantly enriched in sepsis patients. Seventeen key genes were identified and used to classify sepsis into two subtypes via WGCNA and consensus clustering. These subtypes exhibited distinct clinical and immune profiles. Seven hub genes (BPI, HGF, HP, LCN2, LTF, MMP8, RETN) showed differential expression between subtypes and may serve as diagnostic biomarkers. MMP8 was identified as a critical regulator in lactate metabolism, with associated miRNAs and transcription factors predicted.

Single-cell analysis revealed altered immune cell compositions and interactions in sepsis patients. Our findings offer novel insights into sepsis pathogenesis and potential therapeutic strategies targeting lactate metabolism and immune regulation. Keywords: sepsis; lactate metabolism; bioinformatics; immune microenvironment

## Beckground

Sepsis is a systemic inflammatory response triggered by infection and is one of the leading causes of death in intensive care units. (1, 2). It can be initiated by various pathogens, with bacterial infections being the most predominant (3-5). The pathological hallmarks of sepsis are characterized by dysregulated systemic inflammation, which results in tissue injury and organ dysfunction. Clinically, patients often present with high fever, tachycardia, and rapid breathing, which may progress to septic shock and multi-organ failure in severe cases(6-9). The progression of sepsis involves aberrant expression patterns of numerous cytokines and inflammatory mediators. Moreover, the initial phase of sepsis is marked by excessive immune activation, wherein immune cells become highly activated and release inflammatory factors. This may subsequently lead to immune suppression, increased lymphocyte apoptosis, and reduced immune function, thereby elevating the risk of secondary infections(10-12). Besides, sepsis involves cellular reprogramming, including metabolic and epigenetic changes. Metabolic reprogramming alters immune cell function, while epigenetic modifications, such as DNA methylation and histone changes, affect gene expression, influencing sepsis progression and immune responses(13, 14). However, the pathogenesis of sepsis remains unclear completely, posing a significant challenge to clinical treatment. Therefore, elucidating the molecular and immunological mechanisms underlying sepsis is paramount for developing novel treatment strategies and enhancing patient survival rates.

During tissue hypoxia or metabolic disturbances, the lactate metabolism pathway becomes activated, thereby enhancing glycolysis and increasing lactate production. This process exerts a significant impact on immune cell functionality and the regulation of inflammatory responses. Lactate modulates immune cell status and function via multiple mechanisms(15). In the context of sepsis, elevated lactate levels are strongly correlated with increased disease severity and a poorer prognosis. Moreover, Lactylation modification became a potential marker for diagnosis and guiding the treatment of sepsis(16-18). Consequently, investigating lactate metabolism is essential for elucidating the underlying mechanisms of sepsis and for identifying novel therapeutic targets.

Our research primarily investigates the lactate metabolism pathway and the immune microenvironment in sepsis, aiming to provide novel insights into its pathogenesis and to refine clinical treatment strategies. We have systematically analyzed the key genes and pathways involved in lactate metabolism during sepsis and have innovatively classified sepsis into two subtypes based on variations in lactate metabolism. Furthermore, we have comprehensively examined the characteristics of immune cells in sepsis, elucidating the interaction networks among these cells and the expression patterns of immune regulatory molecules. By integrating data on lactate metabolism and immune regulation, we seek to identify novel therapeutic targets and explore immunotherapeutic strategies based on the modulation of lactate metabolism, with the ultimate goal of improving clinical outcomes for patients with sepsis.

## Methods

### 2.1. Data Collection and Download

The dataset GSE134347 comprises peripheral blood samples from 156 sepsis patients, 59 non-infectious ICU patients, and 83 healthy individuals. The dataset GSE65682 includes peripheral blood samples from 192 sepsis patients, 33 non-infectious ICU patients, and 42 healthy individuals. These two datasets were employed to screen for differentially expressed NRGs. All data utilized in this study are publicly available via the GEO database (http://www.ncbi.nlm.nih.gov/geo/). Detailed information regarding the datasets is provided in Table 1.

**Table 1.**

| Accession | Platform | Sample Type | Normal | Non | Sepsis | Gene | Contributors |
| --- | --- | --- | --- | --- | --- | --- | --- |
| GSE134347 | HTA 2.0 | Peripheral blood | 83 | 59 | 156 | mRNA | Brendon Scicluna |
| GSE65682 | HG-U219 | Peripheral blood | 42 | 33 | 192 | mRNA | Brendon Scicluna |

### 2.2. Identification of Differentially Expressed Genes (DEGs)

The R package limma was used to analyze differential expression data, applying the criteria |logFC| > 0.5 and p < 0.05. Volcano plots, heatmaps, and boxplots were generated using the ggplot2 and pheatmap packages.

### 2.3. Enrichment Analysis

KEGG enrichment analysis and Gene Ontology (GO) analysis were conducted separately for the DEGs. Specifically, we utilized the R packages clusterProfiler and GOplot to perform GO functional enrichment analysis (including biological process, cellular component, and molecular function categories) and KEGG pathway analysis. Gene Set Variation Analysis (GSVA) was used to assess changes in pathway activity under specific biological conditions by estimating the relative enrichment of a set of genes within the sample population. Probe ID conversion was performed using the org.Hs.eg.db package, and Z-scores were calculated in conjunction with expression levels via the GOplot package; a Z-score greater than 0 indicates positive regulation, whereas a score less than 0 indicates negative regulation.

### 2.4. Protein–Protein Interaction (PPI) Analysis and Hub Gene Identification

We analyzed the interactions among 17 overlapping genes using the online STRING database and Cytoscape software.

### 2.5. Subtype Analysis Using Consensus Clustering

Based on the differentially expressed genes (DEGs), the ConsensusClusterPlus package in R was employed to perform unsupervised clustering of sepsis samples. Samples were grouped into k = [2, 3, 4, 5, 6] clusters to determine the k value that provided stable clustering. The optimal k was validated using the proportion of ambiguous clustering (PAC).

### 2.6. Relationship Between Subtypes and Clinicopathological Features

Clinical data for each subtype were analyzed using chi-square tests to assess the association between subtypes and various clinicopathological features, including gender, endotype, and diabetes status.

### 2.7. Evaluation of Immune Cells

CIBERSORT was widely utilized to estimate the abundance of immune cells in the tumor microenvironment. The LM22 signature matrix was downloaded from the CIBERSORTx website (https://cibersortx.stanford.edu/). In this study, we employed a deconvolution method based on CIBERSORT along with the LM22 signature matrix to quantify the relative proportions of 22 immune cell subsets in samples from different subtypes. The composition of infiltrating immune cells in each sample was visualized using the *ggplot2* package, and the *ggpubr* and *cowplot* packages were used to compare the 22 immune cell subtypes between the sepsis subtypes. The correlations between different immune cells were analyzed using the *corrplot* package, and Spearman correlation analysis between hub genes and immune cells was conducted using the *ggstatsplot* package.

### 2.8. Receiver Operating Characteristic (ROC) Curve Analysis

ROC curve analysis was performed using the *pROC* and *ggplot2* packages to calculate the area under the curve (AUC) for evaluating the diagnostic efficacy of the hub genes. The AUC, which incorporates both sensitivity and specificity, was used to validate the inherent diagnostic performance of the biomarkers; an AUC greater than 0.5 indicates diagnostic effectiveness, with values closer to 1 reflecting superior performance. In this study, an AUC greater than 0.7 was considered indicative of ideal diagnostic value.

### 2.9. Construction of a TF–miRNA Coregulatory Network

The RegNetwork database (https://regnetworkweb.org) was employed to identify miRNAs and regulatory transcription factors (TFs) associated with the genes of interest. Using this database, TF–miRNA coregulatory interactions were collected for the validated necrosis-related hub genes. Subsequently, the TF–miRNA coregulatory network was visualized using NetworkAnalyst (https://www.networkanalyst.ca), which aids researchers in identifying biological features and functions within complex datasets, thereby forming robust biological hypotheses. The network reflects the interactions between miRNAs and TFs with shared hub gene targets, which may elucidate the regulation of NRG expression.

### 2.10. Analysis of DRGs and Drug Interactions

The drug–gene interaction database DGIdb (https://www.dgidb.org) was used to identify potential therapeutic drugs for sepsis. Specifically, we uploaded 17 hub genes to the DGIdb database to identify potential drugs and molecular compounds that may be associated with these hub genes.

### 2.11. Processing and Analysis of scRNA-seq Data

In this study, raw single-cell RNA sequencing data were subjected to quality control using the R package *Seurat*. Specifically, cells were excluded if mitochondrial content exceeded 20%, if the expression levels of red blood cell-related genes (HBA1, HBA2, HBB, HBD, HBE1, HBG1, HBG2, HBM, HBQ1, and HBZ) exceeded 5%, or if the number of features (nFeature_RNA) was below 200 or above 7500. Additionally, cells with nFeature_RNA counts outside the range of 1 to 100,000 were removed. Only cells meeting these quality control criteria were retained. Following normalization, principal component analysis (PCA) was performed on the top 2,000 highly variable genes. Cell clusters were identified based on the first 20 principal components with a resolution of 0.8. The *SingleR* package was used to assign specific labels to each cluster. Differentially expressed genes (DEGs) for distinct cell populations were identified using the FindAllMarkers function in the *Seurat* package. Finally, cell–cell communication was inferred using the R package *CellChat*, which calculates communication networks (in terms of number and strength) by computing connectivity and communication probabilities between different cell populations.

## Result

### 3.1 Differential Gene Expression and Enrichment Analysis in Sepsis

Two Gene Expression Omnibus (GEO) datasets (GSE134347 and GSE65682) were collected, which include transcriptomic data from healthy individuals, critically ill sepsis patients, and critically ill non-infectious patients. First, differential expression analysis (DEGs) was performed between healthy individuals and sepsis patients, as well as between healthy individuals and non-infectious patients, for both datasets (Figures 1.A-D). Gene Ontology (GO) pathway enrichment analysis revealed that the comparisons between sepsis and healthy controls, as well as between sepsis and non-infectious patients, yielded consistent DEG profiles and GO enrichment patterns across both datasets, involving multiple pathways related to immune response and cellular regulation (Figure 1.E). We further focused on analyzing the role of the lactate metabolism pathway in sepsis. Gene Set Enrichment Analysis (GSEA) demonstrated that, compared with healthy individuals or non-infectious patients, sepsis patients exhibited significant enrichment of the lactate metabolism pathway (Figures 1.F-I).

**Figure 1:**
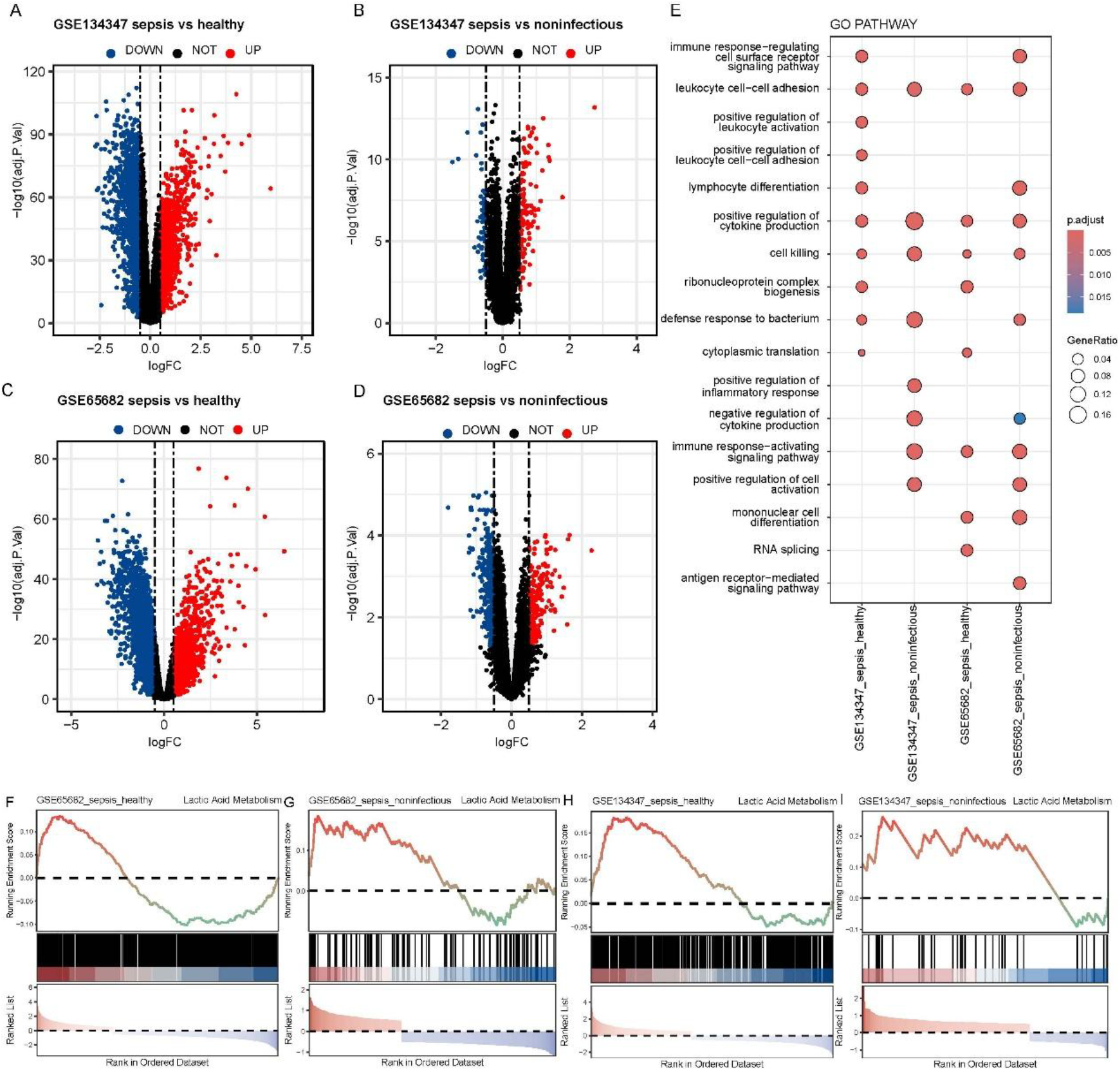
Differential Gene Expression and Pathway Enrichment Analysis. A-D: Volcano plots display the expression profiles of differentially expressed genes (DEGs) between the sepsis group and the healthy or non-infectious groups in the GSE134347 and GSE65682 datasets. E: Bar charts illustrate the GO enrichment analysis for immune response and cellular regulation pathways across the different group comparisons. F-I: GSEA enrichment analysis of the lactate metabolism pathway in the various groups.

### 3.2. Differences in the Lactate Metabolism Pathway Between Sepsis and Healthy Individuals

A Venn diagram (Figure 2.A) illustrates that 17 genes overlap between lactate metabolism-related genes and the DEGs identified from the sepsis versus healthy or non-infectious comparisons in the two datasets. We calculated lactate metabolism pathway scores for these 17 genes using ssGSEA, and observed that sepsis patients exhibited significantly higher pathway scores compared to healthy individuals and non-infectious patients (Figures 2.B and 2.C). Further, heatmap analysis of these 17 overlapping DEGs revealed similar patterns of upregulation and downregulation in both the GSE134347 and GSE65682 datasets; notably, genes such as CD2, TREM1, and MME were downregulated in a subset of sepsis samples (Figures 2.D and 2.E). Subsequently, we constructed a protein–protein interaction (PPI) network based on the proteins expressed by these 17 overlapping DEGs (Figure 2.F), which revealed interactions among proteins encoded by nine genes—MMP8, BPI, LCN2, LTF, RETN, HP, HGF, TREM1, and ALPL. These interactions suggest that a potential regulatory mechanism exists among these genes, implying that the aberrant lactate metabolism pathway observed in sepsis relative to healthy and non-infectious patients may be centered on the regulatory networks of these molecules. To further elucidate the functional mechanisms of these 17 overlapping genes, we identified hub genes within their expression network and performed GO enrichment analysis, which showed the most significant enrichment in the “Defense response to bacterium” category (Figure 2.F).

**Figure 2:**
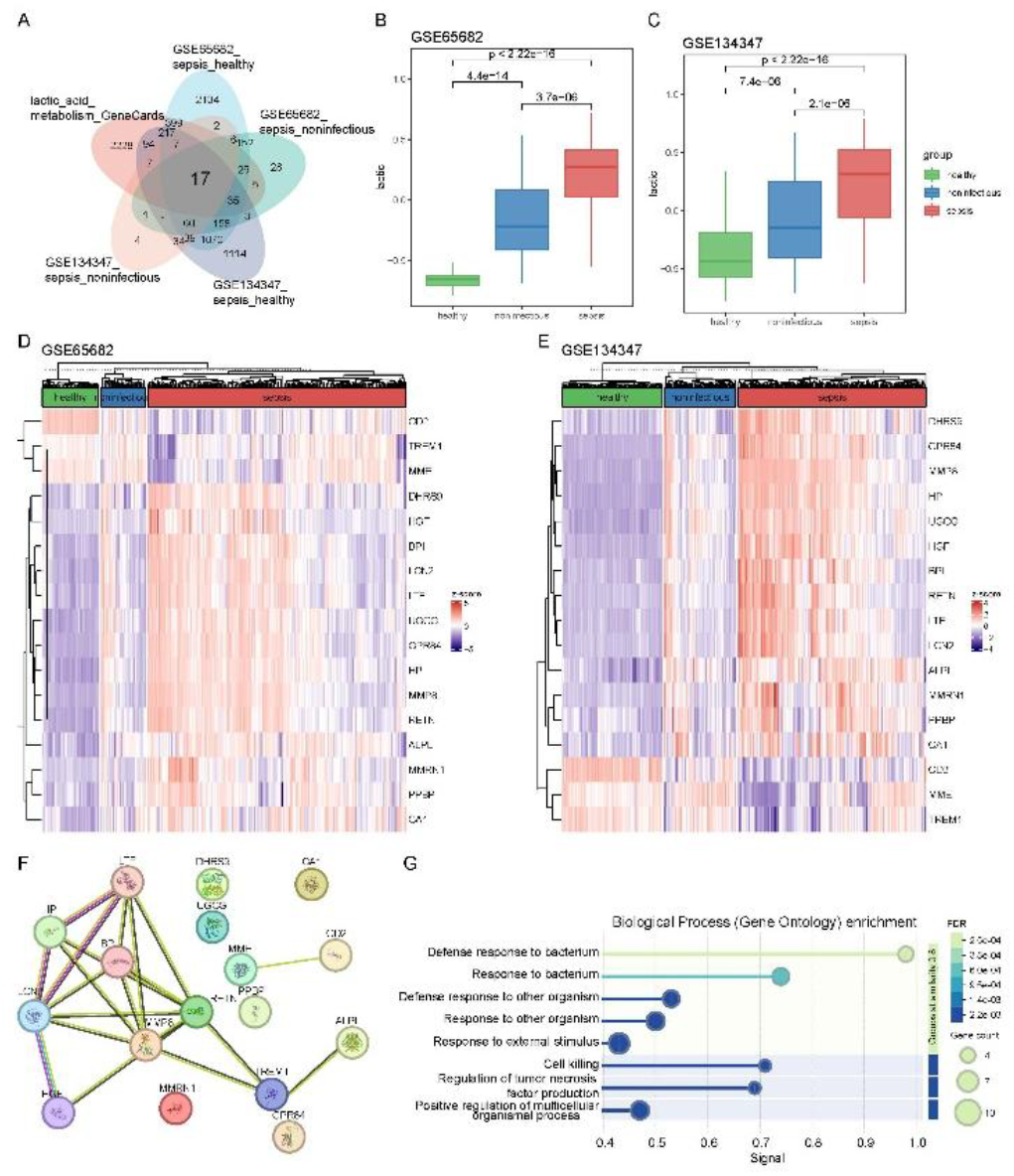
Analysis of the Lactate Metabolism Pathway. A: Venn diagram displaying the overlapping genes. B, C: ssGSEA analysis of lactate metabolism pathway scores; boxplots illustrate the score differences among groups. D, E: Heatmaps showing the expression profiles of the 17 overlapping lactate metabolism-related genes across samples from each group. F: Protein–protein interaction network of the proteins expressed by the 17 overlapping lactate metabolism-related genes. G: GO enrichment analysis of the hub genes within the 17 overlapping lactate metabolism-related genes.

### 3.3. Subtyping Sepsis Patients Based on the Sepsis Gene Set

We conducted an in-depth analysis of the expression data of the hub genes from the lactate metabolism pathway using weighted gene co-expression network analysis (WGCNA) to identify sepsis-associated gene modules and their relationships with clinical traits. Combined analysis of the cumulative distribution function (CDF) curve from consensus clustering (Figure 3.A) and the proportion of ambiguous clustering (PAC) (Figure 3.B) determined an optimal k value of 2. Consequently, unsupervised consensus clustering was applied to classify the sepsis samples into two subtypes, designated as cluster1 and cluster2 (Figure 3.C). Further examination of the clinical features revealed that these subtypes were associated with internal phenotype classifications and the presence of diabetes, but not with gender (Figure 3.D). To explore differences in immune cell populations between the two subtypes, the CIBERSORT algorithm was used to quantitatively assess the abundance of 22 immune cell types in the cluster1 and cluster2 samples. In cluster1, the abundances of immune cells such as Eosinophils, resting Dendritic cells, Macrophages M0, and Monocytes were significantly higher compared to cluster2, whereas the abundances of Neutrophils, resting NK cells, activated CD4 memory T cells, naive CD4 T cells, CD8 T cells, and Tregs were lower (Figure 3.F). Interestingly, the EBV infection pathway exhibited the highest degree of enrichment among the differential pathways between the two sepsis subtypes (Figure 3.E).

**Figure 3:**
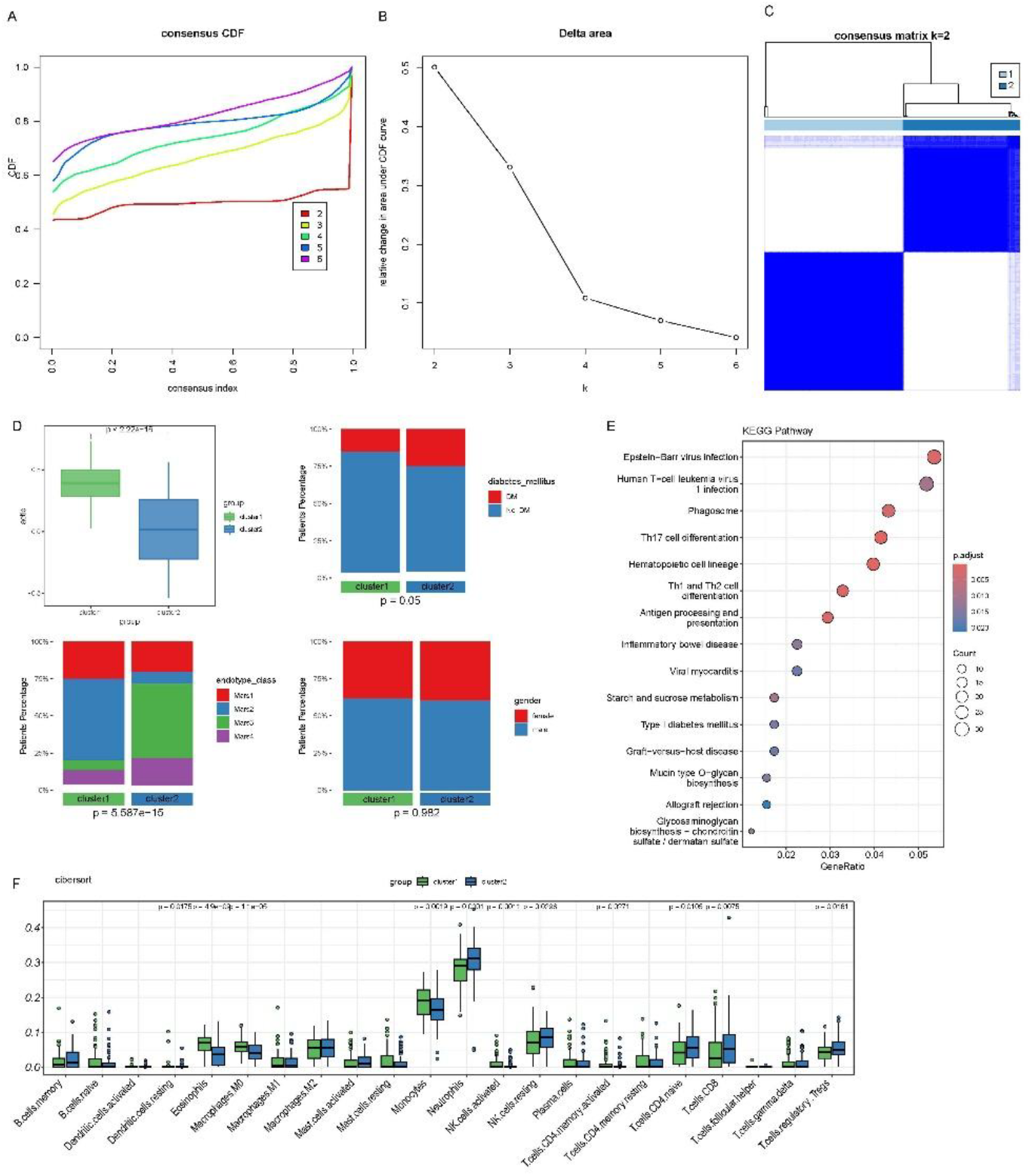
Identification and Characterization of Subtypes. A. Cumulative distribution function (CDF) curve for consensus clustering. B. Proportion of ambiguous clustering (PAC) analysis. C. Heatmap of subtype clustering. D. Bar charts and boxplots displaying the clinical characteristics of cluster1 and cluster2 populations. E. KEGG pathway enrichment analysis of differential genes between the two subtypes. F. Comparison of the levels of 22 immune cell types between the two subtypes.

### 3.4. Clinical Subtyping Markers for the Two Subtypes

Further analysis of the 9 potential interacting hub genes in the two subtypes revealed that the expression levels of 7 genes—BPI, HGF, HP, LCN2, LTF, MMP8, and RETN—were elevated in the cluster1 subtype and reduced in the cluster2 subtype (Figure 4.A). Analysis of 22 immune cell types also demonstrated divergent expression trends for these 7 hub genes between the two subtypes; specifically, both the GSE134347 and GSE65682 datasets showed a significant increase in Macrophages M0 and Eosinophils, while Neutrophils exhibited a significant decrease (Figures 4.B and 4.C). Moreover, the ROC curves for the diagnostic performance of each characteristic gene in both datasets displayed a good fit. These findings suggest that these 7 hub genes may serve as diagnostic markers for distinguishing different clinical subtypes of sepsis, thereby providing potential targets for diagnosis or therapeutic intervention (Figures 4.D–G).

**Figure 4:**
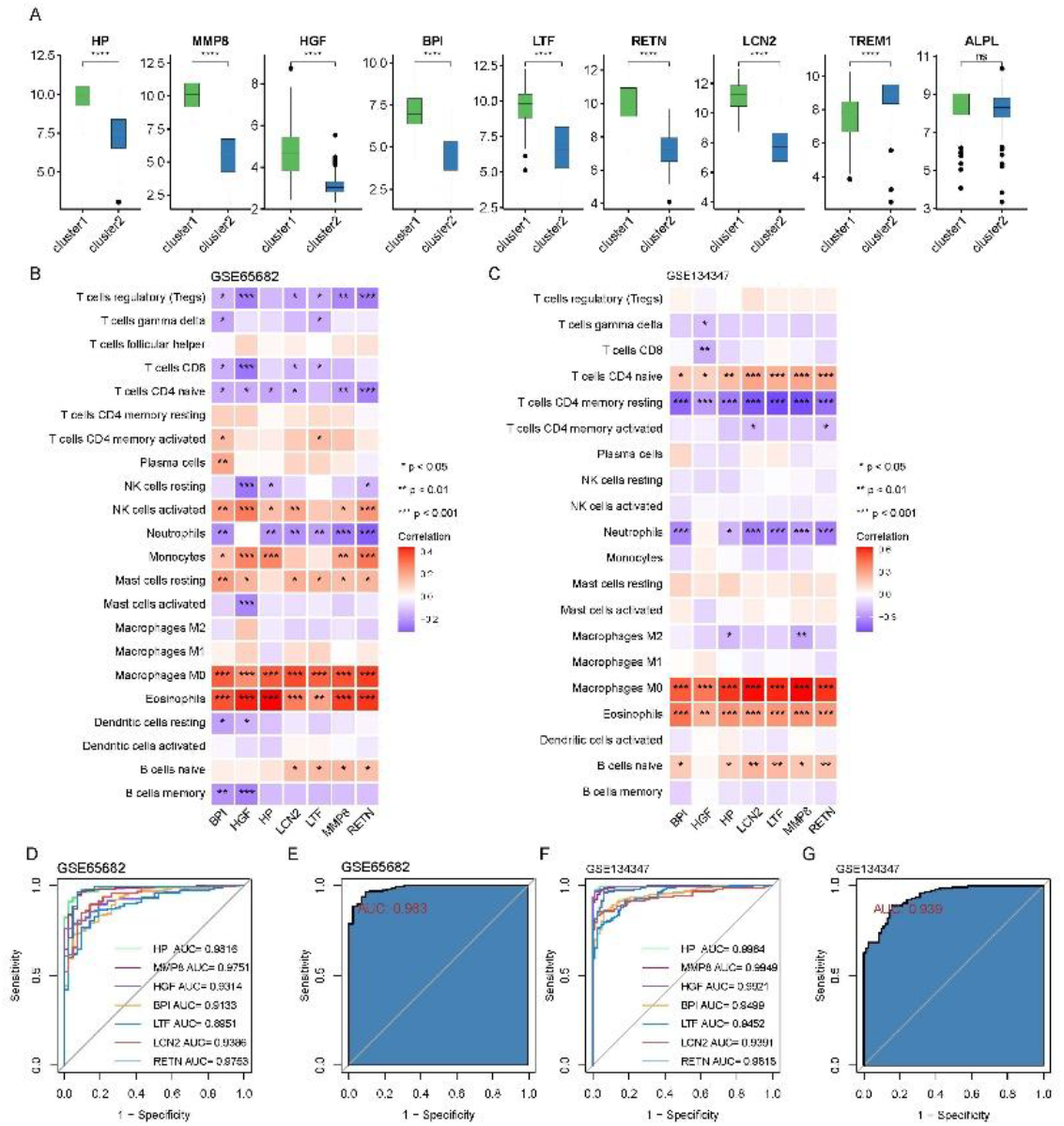
Clinical Subtyping Markers in the Two Subtypes. A: Boxplots illustrating the expression differences of 9 hub genes between the two clusters. B, C: Correlation analysis between the hub genes and 22 immune cell types in both datasets. D, F: ROC curves showing the diagnostic performance of each hub gene in both datasets. E, G: ROC curves depicting the integrated diagnostic performance of the 7 hub genes in both datasets.

### 3.5. MMP8 Gene-Related Analysis

Previous findings revealed that the MMP8 gene may play a key regulatory role within the lactate metabolism pathway in sepsis (Figure 2.F). To uncover the potential mechanisms underlying MMP8 expression in sepsis, we classified samples from both datasets into groups with high and low MMP8 expression. Through GSVA and KEGG pathway analysis, we observed that multiple pathways associated with immune cell responses were significantly enriched in samples with differing MMP8 expression levels in both the GSE134347 and GSE65682 datasets (Figures 5.A–D). Furthermore, we predicted several miRNAs that may be involved in the regulation of MMP8 in sepsis (Figure 5.E). In addition, transcription factor prediction using various databases consistently identified STAT1 as a potential regulator of MMP8 expression (Figure 5.F). These findings suggest that the differential expression of MMP8 in sepsis patients may be linked to distinct molecular mechanisms involving multiple pathways and regulatory molecules.

**Figure 5:**
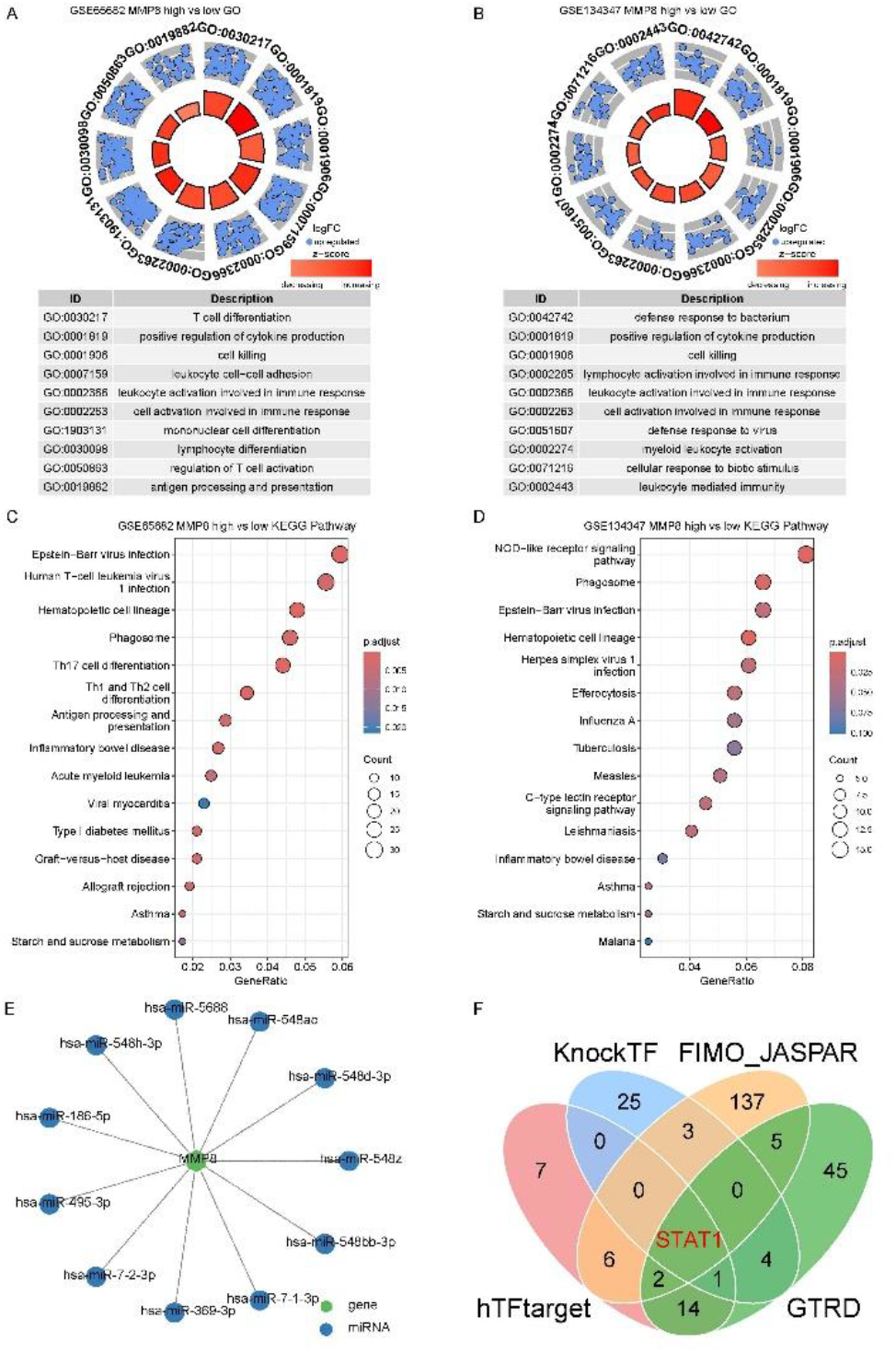
MMP8 as a Key Differential Regulatory Gene in Sepsis. A, B: GSVA scores of HALLMARK pathways that are significantly different between high and low MMP8 expression samples. C, D: Enrichment scores for KEGG pathways that are significantly different between high and low MMP8 expression samples. E: Predicted miRNAs interacting with the MMP8 transcript. F: Predicted transcription factors regulating MMP8 expression.

**Figure 6:**
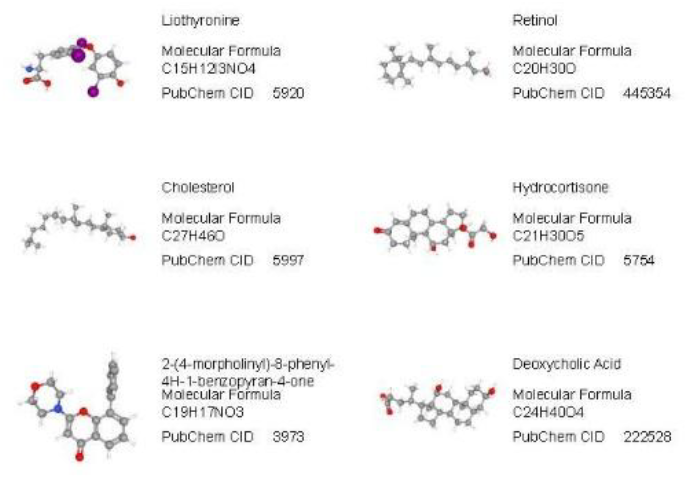
Prediction of potential small-molecule drugs for treating sepsis based on the 7 hub genes.

**Figure 7:**
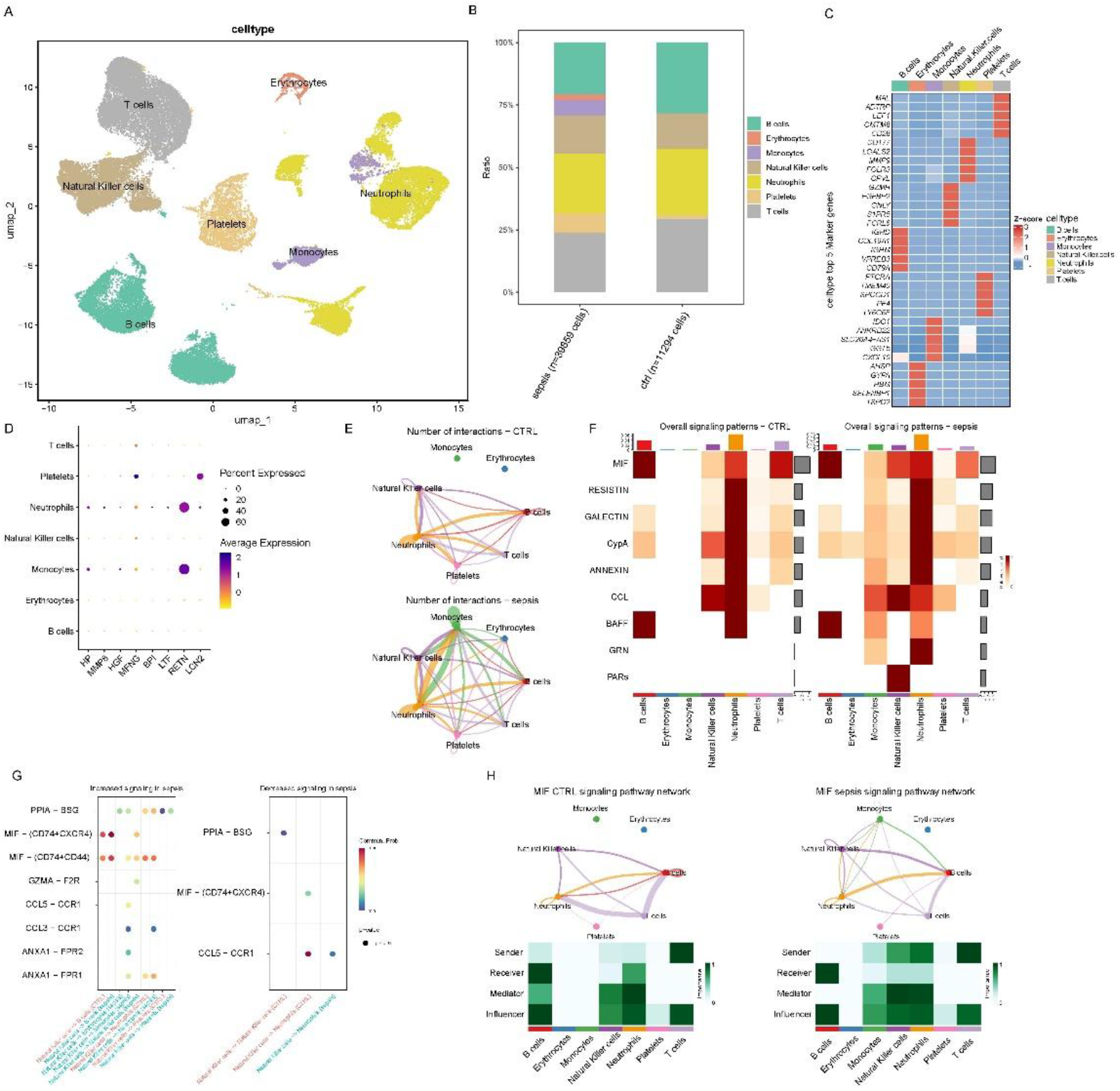
Single-cell transcriptomic data reveals cellular communication processes in sepsis A: UMAP plot clusters seven cell types. B: Stacked plot shows the proportion of each cell type. C: Heatmap displays the top five differentially expressed genes in each cell type. D: Dot plot shows the expression levels of hub genes in each cell type. E: Interaction network plot among different cell types. F: Heatmap shows signal molecules involved in regulating cell communication. G: Bubble plot shows differential cell communication between CTRL and sepsis groups. H: Interaction network plot of the MIF signaling pathway in different cell types and a heatmap showing MIF signaling pathway activity.

### 3.6. Molecular Identification of Candidate Drugs

We queried the DSigDB database to identify potential drug compounds that may interact with the 7 hub genes associated with the lactate metabolism pathway. Based on the integrated scores, the top six predicted potential drug compounds are as follows: **Liothyronine**; **Retinol**; **Cholesterol**; **Hydrocortisone**; **2-(4-Morpholinyl)-8-phenyl-4H-1-benzopyran-4-one; Deoxycholic Acid**. These compounds have been identified as potential therapeutic agents targeting the key regulatory genes involved in the lactate metabolism pathway associated with sepsis. Further experimental validation is necessary to assess their efficacy and safety in clinical settings.

### 3.7. Single-cell transcriptomic data analysis results

Here, we performed dimensionality reduction, clustering, and annotation on scRNA-seq data from sepsis. Cell populations were categorized into seven types: NK cells, T cells, monocytes, B cells, erythrocytes, platelets, and neutrophils. Compared to controls, sepsis patients had higher proportions of NK cells, monocytes, erythrocytes, and platelets. The RETN gene, one of seven hub genes in the lactate metabolism pathway, was highly expressed in neutrophils and monocytes. Communication analysis revealed strong interactions between monocytes and other cell populations in sepsis. Our analysis of signaling molecules showed that macrophage migration inhibitory factor (MIF) played a significant role in cell communication regulation in sepsis. Further investigation of MIF-related signaling pathways indicated that the MIF gene was involved in monocyte communication in sepsis samples but not in controls. These findings offer initial insights into the immune cell status and communication mechanisms in sepsis patients.

## Discussion

Sepsis is a leading cause of death in ICUs, posing significant therapeutic challenges due to its high incidence and mortality rates. Elucidating its molecular and immune mechanisms is crucial for developing new treatment strategies and improving patient survival. This study employed bioinformatics analysis to explore sepsis pathogenesis and identify potential diagnostic and therapeutic targets.

We analyzed transcriptomic data from two GEO datasets (GSE134347 and GSE65682) encompassing healthy individuals, sepsis patients, and non - infected critically ill patients. Through differential expression analysis and GSEA, we found that the lactate metabolism pathway was significantly enriched in sepsis patients compared to healthy individuals or non - infected patients. Further analysis of the overlap between lactate metabolism - related genes and DEGs identified 17 key genes. We then constructed a PPI network and screened out nine potentially interacting genes that may play important roles in sepsis. Our analysis revealed that the enriched genes are associated with the Defense response to bacterium pathway, a finding consistent with previous studies(19). This suggests their potential role in the immune response to bacterial infections in sepsis.

Using WGCNA and consensus clustering, we classified sepsis samples into two subtypes (cluster1 and cluster2) and analyzed their clinical characteristics and immune cell composition. In cluster1, the abundance of eosinophils, resting dendritic cells, M0 macrophages, and monocytes was elevated, while neutrophils, resting NK cells, and CD4+ memory - activated T cells were reduced. Many cells related to sepsis have been studied(20-22), but there is no specific research on their expression differences and classification of related clinical subtypes We analyzed nine hub genes in the two subtypes and found that BPI, HGF, HP, LCN2, LTF, MMP8, and RETN were up - regulated in cluster1 and down - regulated in cluster2. These genes have been sporadically reported in relation to sepsis, but there’s no comprehensive combined analysis(23-26). We innovatively link these hub genes as one clinical subtype. These genes may serve as diagnostic biomarkers for sepsis subtypes.

ROC curve analysis confirmed the diagnostic efficacy of these seven hub genes for sepsis subtypes, offering potential targets for precision medicine. GSVA and GEGG pathway analysis indicated that MMP8 might play a key regulatory role in sepsis - associated lactate metabolism. Studies have explored the link between MMP genes and sepsis(27). MMP8 is a shared biomarker for T2DM and sepsis(28). Our sepsis subtypes are associated with diabetes, and further research is needed to understand the connection between this and lactate metabolism regulation. We predicted miRNAs and transcription factors involved in MMP8 regulation and identified six potential drug compounds targeting the seven hub genes in the DSigDB database. These drugs could provide new therapeutic strategies by modulating key genes in the lactate metabolism pathway.

Single - cell transcriptomic analysis revealed increased NK cells, monocytes, erythrocytes, and platelets in sepsis patients. RETN was highly expressed in neutrophils and monocytes. Monocytes had strong interactions with other cell populations, and MIF played a significant role in cell communication regulation in sepsis.

However, this study has limitations. It only preliminarily explored the lactate metabolism pathway and immune cell status in sepsis patients. Further research is needed to understand the complex regulatory mechanisms and interactions between lactate metabolism and immune cells, as well as to analyze the expression differences of the lactate metabolism pathway between the two subtypes. Additionally, more comprehensive studies are required to investigate monocytes in sepsis, as our analysis was based on a single - cell dataset. Future research should address these aspects to deepen our understanding of sepsis pathogenesis and develop more effective therapeutic strategies.

## Conclusion

Our study revealed that the lactate metabolism pathway is enriched in sepsis patients. We identified key genes and classified sepsis into two subtypes with distinct immune cell profiles. These findings offer new insights into sepsis pathogenesis and potential targets for therapy.

## Notes

### Competing Interest Statement

The authors have declared no competing interest.

